# Liver stage fate determination in *Plasmodium vivax* parasites: characterization of schizont growth and hypnozoite fating from patient isolates

**DOI:** 10.1101/2022.06.16.496373

**Authors:** Amélie Vantaux, Julie Péneau, Caitlin A. Cooper, Dennis E. Kyle, Benoit Witkowski, Steven P. Maher

## Abstract

*Plasmodium vivax*, one species parasite causing human malaria, forms a dormant liver stage, termed the hypnozoite, which activate weeks, months or years after the primary infection, causing relapse episodes. Relapses significantly contribute to the vivax malaria burden and are only killed with drugs of the 8-aminoquinolone class, which are contraindicated in many vulnerable populations. Development of new therapies targeting hypnozoites is hindered, in part, by the lack of robust methods to continuously culture and characterize this parasite. As a result, the determinants of relapse periodicity and the molecular processes that drive hypnozoite formation, persistence, and activation are largely unknown. While previous reports have described vastly different liver stage growth metrics attributable to which hepatocyte donor lot is used to initiate culture, a comprehensive assessment of how different *P. vivax* patient isolates behave in the same donors at the same time is logistically challenging. Using our primary human hepatocyte-based *P. vivax* liver stage culture platform, we aimed to simultaneously test the effects of how hepatocyte donor and *P. vivax* patient isolate influence the fate of sporozoites and growth of liver schizonts. We found that, while environmental factors such as hepatocyte donor can modulate hypnozoite formation rate, the *P. vivax* case is also an important determinant of the proportion of hypnozoites observed in culture. In addition, we found schizont growth to be mostly influenced by hepatocyte donor. These results suggest that, while host hepatocytes harbor characteristics making them more-or less-supportive of a quiescent versus growing intracellular parasite, sporozoite fating towards hypnozoites is isolate-specific. Future studies involving these host-parasite interactions, including characterization of individual *P. vivax* strains, should consider the impact of culture conditions on hypnozoite formation, in order to better understand this important part of the parasite’s lifecycle.

**Author summary:** Malaria is caused by protozoan parasites of the genus *Plasmodium*. One species, *Plasmodium vivax*, is more difficult to control in comparison to other species because infection results in dormant forms in the liver, called hypnozoites. Hypnozoites are considered an invaluable therapeutic target to control malaria, but how hypnozoites form and reactive to cause malaria relapses is unknown. Herein we describe that both nature and nurture influence the fate of newly-established parasites in the liver, resulting in either a quiescent hypnozoite or growing schizont. Using parasites generated from patient isolates, we show the hypnozoite formation is likely inherited but also modulated by environmental factors, including which lot of human hepatocytes the parasites infect. Additionally, we show schizont growth is strongly influenced by the host hepatocyte lot. As liver stage experiments include several dependent variables which are difficult to control, herein we present an experimental approach designed to remove many of these variables and provide a clearer picture of what factors influence the formation and growth of liver stage parasites. Our findings serve as a foundation for future work to understand hypnozoite biology, with the ultimate goal of identifying new therapeutic targets.

## Introduction

Malaria remains a major public health challenge with an estimated 241 million cases estimated in 2021 [1]. Among the parasite species responsible for human malaria, *Plasmodium vivax* is the most widely dispersed as well as the most resistant to elimination programs. This resilience is attributed to several features unique to *P. vivax*, including its ability to develop over a wider range of temperatures and, in particular, at lower thermal limits than *Plasmodium falciparum*. Additionally, *P. vivax* forms transmissible gametocytes faster, and has a shorter incubation period in the mosquito vector, than *P. falciparum* [2-5]. Critically, *P. vivax* parasites persist in the human host liver as hypnozoites (a dormant parasite liver stage) which activate weeks, months or years after the primary infection, causing relapse episodes [6]. Hypnozoites are insensitive to most antimalarials except 8-aminoquinolines, which are contraindicated in large sections of the population including pregnant women, younger children, and patients with glucose-6-phosphate dehydrogenase deficiency [7]. Because *in vitro* culture of the liver-stages is dependent on limited access to *P. vivax* infected mosquitoes, our understanding of *P. vivax* liver-stages has considerably lagged in comparison to other *Plasmodium* species. Consequently, the determinants of relapse periodicity and the molecular processes that drive hypnozoite formation, persistence, and activation are still largely unknown.

Transmission occurs when sporozoites are injected into a new host by the bite of an infected *Anopheles* mosquito. Individual sporozoites migrate to the liver, invade a hepatocyte and form either a liver-schizont or hypnozoite. Schizonts mature within 9-12 days and release merozoites into the bloodstream, thereby initiating the primary blood stage infection, while hypnozoites are small, non-dividing forms that remain quiescent for various periods of time [8]. Frequencies of *P*.*vivax* relapses are highly variable, from 3-4 weeks in the tropics to 8-10 months in temperate regions, and it remains unknown if the frequencies observed are determined by genetic or environmental factors [6, 8-10]. Several factors influencing relapses frequencies have been proposed such as sporozoites inoculum size, acquired immunity of the host, primary drug treatment regimens, co-infections, fever, hemolysis, seasonality, mosquito bites, and epigenetic control [6, 9, 11-13]. However, a constant activation rate without external stimuli could also explain the frequencies observed [10]. Yet, the prevalence of hypnozoite formation is rarely considered and difficult to directly ascertain in living systems [14, 15].

The cellular interactions governing migration and invasion of sporozoites into hepatocytes are species-specific and only partially understood, rendering generalizations difficult [16, 17]. Comparisons of infection rates in several human hepatocyte donors show that some donors are not supportive of either *P. vivax* or *P. falciparum* parasites, which could be a product of natural variation in hepatic surface receptors necessary to malaria parasite entry [17]. Alternatively, the process for manufacturing cryo-plateable lots of primary hepatocytes could affect cell phenotypes, including the aforementioned surface receptors, and alter hepatocyte permissiveness [18, 19]. Host cell permissiveness is likely also modulated by host cell environment as sufficient glycolytic and respiratory activities are needed to sustain the energy demands of an intracellular parasite [20-22]. As such, liver lobules perform different metabolic functions and have recently been shown to influence *P. falciparum* parasite preferences and growth in the host cell [22]. Interestingly, a recent single-cell transcriptomic study of *P. vivax* liver stages did not show a clear pattern of infection in different hepatocyte subpopulations, although, it is unknown if zonally-differentiated hepatocytes remain fully differentiated *ex vivo* [23]. Thus, human host hepatocyte characteristics are likely important factors in parasite development as well as potential determinants of the schizont or hypnozoite fate remaining to be discovered.

The density of individuals within a shared environment has a strong impact on individual fitness. For parasites such as *Plasmodium* spp., fitness depends on interactions with several organisms during their life cycle, including but not limited to, the human host, the mosquito vector, and co-infecting malaria parasites. The host represent ecological niches for co –infecting malaria parasites, which often consist of more than one parasite genotype [24, 25]. Therefore, individual parasites are in direct competition for resources, in indirect competition with shared exposure to immune responses, and potentially in direct interference between parasites, which can all affect virulence and transmission [24-27]. Although much work has been carried out on asexual stages due to the availability of culture and analyses, these bottom-up and top-down mechanisms could also affect sporozoite fate akin to how crowding and inbreeding rate, among other factors, influence gametocyte fate and infectivity [5, 28, 29]. Thus, during transmission from the vector to the human host, parasite competition could favor the production of hypnozoite-fated sporozoites to decrease future competition in the human host and increase the likelihood of relapse during the next high-transmission season.

Using our recently-developed primary human hepatocytes (PHH)-based 384-well *P. vivax* liver stage culture platform [18, 19], we aimed at testing the effects of hepatocyte donors and *P. vivax* cases on liver-stage parasites. While we have previously reported how sporozoites behave when infected into different donors in this system, due to logistical challenges these studies relied on historical comparisons of sporozoites from only a single *P. vivax* isolate for each experimental run [18]. This original approach cannot account for factors which are known to or likely affect parasite viability and phenotypes, such as the effect of an international shipment needed to send infected mosquitoes from an endemic area to a research laboratory, the health and genetic drift of a mosquito colony over time, the variation of different seedings of each lot of hepatocytes, specific lots of reagents like culture media, the conditions of the laboratory environment and equipment, and which human operators perform dissection, infection, and media replacement during culture. To effectively remove or better control for these factors, this assessment relied on an experimental design in which the same four human donor lots, which were pre-validated to support *P. vivax* infection, were seeded into different wells of the same microtiter plates on the same day and infected with the same inoculum of sporozoites from three different *P. vivax* patient isolates. We used this design to confirm that hepatocyte donors influence the total number of parasites and investigating if hepatocyte donors affected schizont growth and the proportion of hypnozoites observed. Additionally, to characterize other factors critical for establishing *in vitro* culture, we factored into our design an assessment of how the sporozoite inoculum size and *P. vivax* case impacted these three aspects (total number of parasites, schizont size and proportion of hypnozoites).

## Material and methods

### Clinical isolates & collection of *P. vivax* sporozoites

Blood samples were collected from symptomatic *P. vivax* patients at local health facilities in Mondulkiri province (eastern Cambodia) from 2018-2021. Clinical isolate collection and research procedures were reviewed and approved by the Cambodian National Ethics Committee for Health Research (approval number: 100NECHR, 113 NECHR, 104 NECHR). The protocols conform to the Helsinki Declaration on ethical principles for medical research involving human subjects (version 2002) and informed written consent was obtained for all volunteers, or their parent or legual guardian for participant under 18 years old. Patients presenting signs of severe malaria, infected with non-vivax malaria parasites, under 5 years of age, pregnant, or lactating were excluded from the collection. Following informed consent from eligible study participants, venous blood samples were collected by venipuncture into heparin-containing tubes. Immediately after collection, treatment was provided by local health staff according to Cambodia National Malaria Treatment Guidelines. Clinical isolates were immediately prepared for feeding to *Anopheles dirus* mosquitoes in a secure insectary as previously described [19]. Following a *P. vivax* gametocyte-containing bloodmeal, *An. dirus* mosquitoes were maintained on a 10% sucrose+0.05% para-aminobenzoic solution. Mosquitoes found positive for *P. vivax* oocysts at six-days post feeding were transported to the IPC facility in Phnom Penh, Cambodia where salivary glands were aseptically dissected into RPMI without sodium bicarbonate on 16-21 days post-infection (dpi).

### Liver stage infection

Primary human hepatocytes (PHH) were seeded 2 days prior to infection (except for experiment 1 for which they were seeded on the same day) and cultured as previously described [19]. Infection was performed by diluting freshly-dissected sporozoites into culture media with antibiotics, adding 20μL sporozoite-media mixture to each well, and centrifugation of the 384-well plate at 200 RCF for 5 min at room temperature. Media was changed with fresh culture media containing antibiotics the day after infection and every 2-3 days thereafter. At 8 or 12 dpi (depending on the experiment protocol, see below) cultures were fixed for 15 min at room temperature with 4% paraformaldehyde in PBS. Fixed cultures were stained with recombinant mouse anti-*P. vivax* Upregulated in Infectious Sporozoites-4 antibody [30] diluted 25,000-fold followed by rabbit anti-mouse Alexfluor488-conjugated antibody diluted 1:1000. Cultures were then counterstained with 10 μg/mL Hoechst 33342 to detect parasite and host cell nuclear DNA. Automated High Content Imaging was carried out with a 20x objective on a ImageXpress Confocal Micro (Molecular Devices) or 4x objective on a Lionheart (Biotek). Liver stage parasite were quantified for number and growth area per well and per parasite using built-in cellular analysis and quantification software (MetaXpress for ImageXpress or Gen5 for Lionheart). Hypnozoites were defined as brightly UIS4-stained round forms (ratio of maximum and minimum widths of each form > 0.6) with under 150 μm^2^ total area and a bright prominence in the PVM. Schizonts were defined as brightly-UIS4-stained forms with greater than 150 μm^2^ total area.

### Experiment 1: testing simultaneously different PHH donors, P. vivax cases and sporozoite inoculum sizes

Four PHH lots, UBV, BGW, HHR and OTW, were infected with three different *P. vivax* cases (C1 to C3) at 8 sporozoite densities ranging from 3 ×10^3^ to 30 ×10^3^ per well in 3 ×10^3^ increments (S1 Fig). Six technical replicate wells per condition were used. Cultures were fixed at 8 dpi.

### Experiment 2: comparison of eight PHH donors and influence of sporozoite inoculum size

Eight PHH lots, BGW, BPB, ERR, HDS, HLY, IRZ, ZPE, were infected with one *P. vivax* case at 6 sporozoite densities ranging from 5 ×10^3^ to 30 ×10^3^ per well in 5 ×10^3^ increments. Four technical replicate wells per conditions were used. Cultures were fixed at 8 dpi.

### Experiment 3: comparison of 51 *P. vivax* cases using one PHH donor

Negative control wells containing 0.1% v/v DMSO from our large *P. vivax* liver stage drug screening database were used to compare results from 51 *P. vivax* cases used to infect 132 assay plates seeded with PHH lot BGW and comprising 106 plates with and 26 plates without 1-aminobenzotriazole (ABT) treatment, a protocol condition used to limit phase I hepatic metabolism of unoptimized test compounds [19]. The number of hypnozoites and schizonts were averaged over 16 or 24 wells DMSO control wells depending on the plate map. The majority of cultures were seeded with PHHs two days prior to infection with sporozoites, although for some infections cultures were initiated one or three days prior to infection due to logistical constraints. All cultures were fixed at 12 dpi.

## Statistical Analyses

### Experiment 1: testing different PHH donors, P. vivax cases and sporozoite inoculum sizes

The total number of parasites was analyzed using Generalized Linear Mixed Model (GLMM) with a zero-truncated negative binomial error structure. In this GLMM, hepatocyte donors (4 levels: BGW, HHR, OTW and UBV), *P. vivax* case (3 levels: C1, C2 and C3) and sporozoite inoculum sizes (12 × 10^3^, 15 × 10^3^, 17 × 10^3^, 19 × 10^3^, 21 × 10^3^, 24 × 10^3^, 27 × 10^3^ and 30 × 10^3^) were coded as fixed categorical factors, the log of number of nuclei was coded as an offset to account for the number of hepatocyte per well, and well nested in plate were coded as random factors to account for repeated measurements of the same infection conditions and plate effects. The proportion of hypnozoites were analyzed using a similar GLMM with a binomial error structure and individual well ID was added as a random factor to improve model fit.

The schizont size (individual measurement of all schizonts present in the well) was Box-Cox transformed and analyzed using a GLMM with a Gaussian distribution. In this model, hepatocyte donors, *P. vivax* case, sporozoite inoculum size and all two-ways interactions were coded as fixed categorical factors and well nested in plate were coded as random factors.

### Experiment 2: comparison of eight PHH donors and influence of sporozoite inoculum size or seed density

The total number of parasites and proportion of hypnozoites were analyzed using GLMMs with a zero-truncated negative binomial error structure and a binomial error structure respectively. In these GLMMs, hepatocyte donors and sporozoite inoculum size were coded as fixed categorical factors, the log of number of nuclei was coded as an offset to account for the number of hepatocytes per well, and replicate wells were coded as random factors to account for repeated measurements of the same infection conditions.

### Experiment 3: comparison of 51 P. vivax cases using one PHH donor

The total number of parasites and proportion of hypnozoites were analyzed using univariate GLMMs with a zero-truncated negative binomial error structure and a binomial error structure respectively. In these GLMMs, the log of the average number of nuclei was coded as an offset to account for the number of hepatocytes per well, and *P. vivax* case was coded as random factor to account for measurements of assay plates infected with the same *P. vivax*. We investigated the effects of sporozoite inoculum size, hepatocyte age at infection, assay version, season and patient sex. In addition, a data subset of 37 plates for which the visit count was known (that is, the number of times the same patient visited the clinic, which was either 2, 3, 5 or 6 visits) was used to investigate the effect of multiple visits on the proportion of hypnozoites using a univariate GLMM with a binomial structure.

Model selection was used with the stepwise removal of terms, followed by likelihood ratio tests (LRT). Term removals that significantly reduced explanatory power (*P*<0.05) were retained in the minimal adequate model [31]. All analyses were performed in R v. 4.0.3 [32]. Results are presented as mean ± standard error (SE) and proportion ± confidence interval (CI).

## Results

### Experiment 1: testing different PHH donors, *P. vivax* cases and sporozoite inoculum sizes

The average number of parasites per well was significantly influenced by the PHH lot (X^2^_3_ = 1023.78, P <0.0001; Fig 1A), with BGW supporting the highest mean parasite per well (260.85 ± 9.64) followed by UBV (162.12 ± 6.09), HHR (123.48 ± 4.74) and OTW (79.28 ± 3.65). The number of parasites was also influenced by the *P. vivax* case (X^2^_2_ = 13.67, P =0.001; Fig 1B) with *P. vivax* case 3 having the highest number of parasites per well (173.95 ± 8.68), followed by *P. vivax* case 1 (168.98 ± 7.64) and *P. vivax* case 2 (126.36 ± 4.77). As expected, the average number of parasites per well increased with the sporozoite inoculum size (X^2^_7_ = 228.37, P <0.0001; Fig 1C), increasing from 89.97 ± 6.76 parasites per well when 12 × 10^3^ sporozoite were inoculated to 211.94 ± 16.66 parasites per well with 30 × 10^3^ sporozoites.

**Fig 1.**
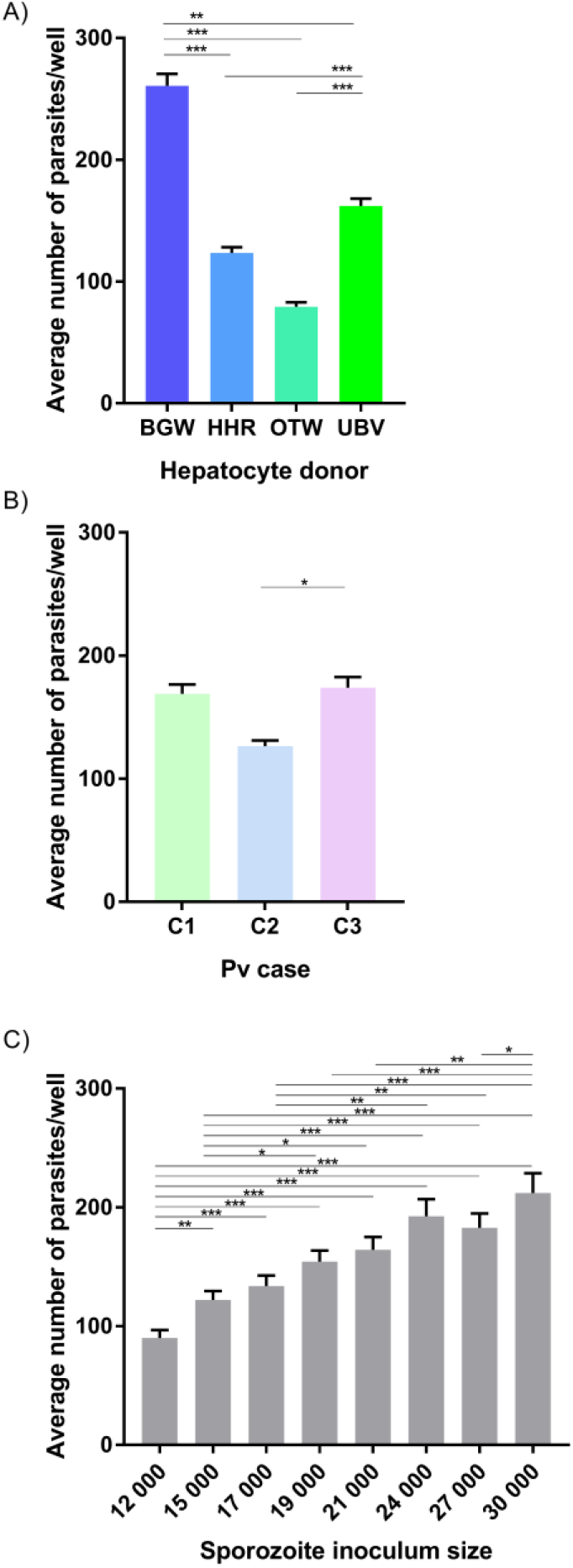
The average number of *P. vivax* parasites per well. Data include all parasites quantified from Experiment 1 and are shown categorized by A) PHH lot, B) *P. vivax* isolate, and C) sporozoite inoculum size. Asteriks indicate significant differences (Post hoc Tukey’s pairwise comparisons, *** P<0.0001, ** P<0.001, * P<0.05). Bars represent ± SE.

The proportion of hypnozoites per well was significantly influenced by the PHH lot (X^2^_3_ = 589.50, P <0.0001; Fig 2A) with UBV supporting the highest proportion (65.11 ± 0.31%) followed by BGW (63.97 ± 0.31%), OTW (54.37 ± 0.32%) and HHR (52.36 ± 0.33%). The three *P. vivax* cases investigated had significantly different proportion of hypnozoites from 57.9 ± 0.32% in *P. vivax* case 1, to 61.8 ± 0.32% in *P. vivax* case 2 and 62.89 ± 0.31% in *P. vivax* case 3 (X^2^_2_ = 50.73, P <0.0001; Fig 2B). The proportion of hypnozoites also showed a small increase as the sporozoite inoculum size increased, from 58.6 ± 0.32% when 12 × 10^3^ sporozoite were inoculated to 63.68 ± 0.31% with 30 × 10^3^ sporozoites (X^2^_7_ = 20.09, P = 0.005; Fig 2C).

**Fig 2.**
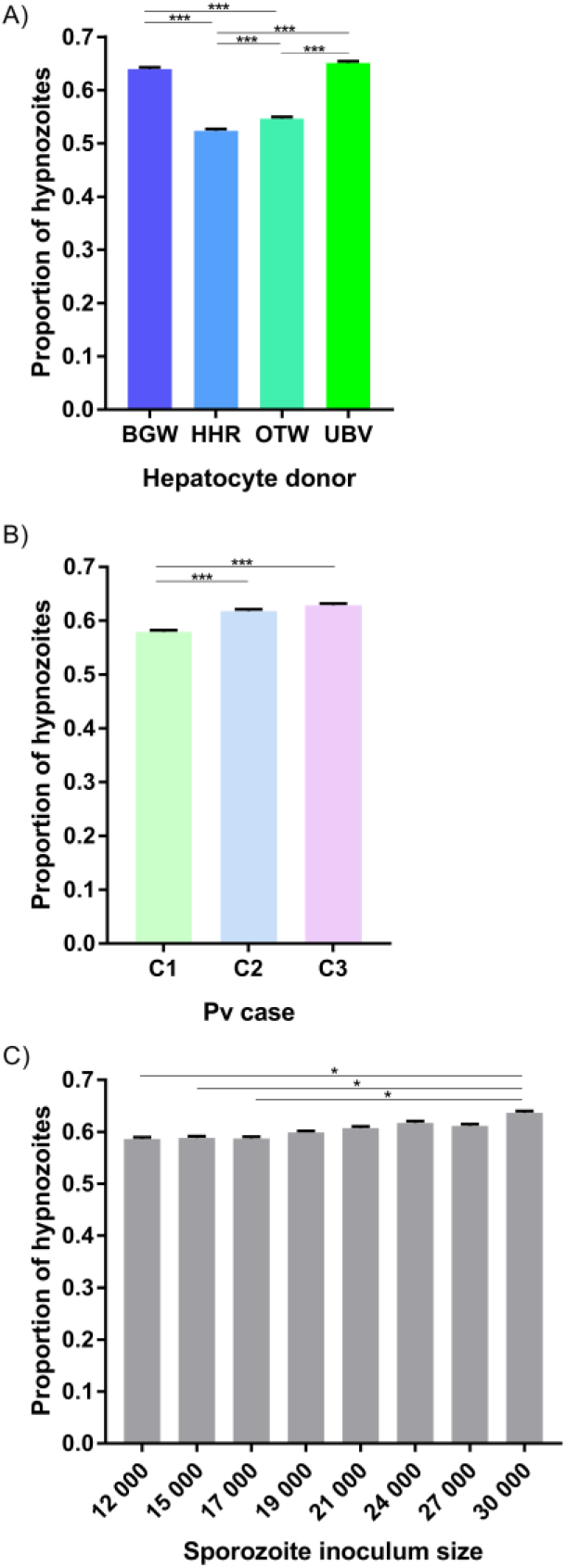
The average proportion of *P. vivax* hypnozoites per well. Data include all parasites quantified from Experiment 1 and are shown categorized by A) PHH lot, B) *P. vivax* isolate, and C) sporozoite inoculum size. Asteriks indicate significant differences (Post hoc Tukey’s pairwise comparisons, *** P<0.0001, ** P<0.001, * P<0.05). Bars represent ± 95% CI.

The schizont size was significantly affected by the PHH lot (X^2^_3_ = 4369.66, P <0.0001; Fig 3A) with an average schizont size of 1195.46 ± 8.25 µm^2^ for HHR, followed by OTW (1021.61 ± 9.16 µm^2^), UBV (443.49 ± 2.94µm^2^) and BGW (441.85 ± 2.83 µm^2^). Schizonts were on average larger in the infection from *P. vivax* case 3 (770.31 ± 6.41µm^2^) followed by *P. vivax* case 1 (678.02 ± 4.81µm^2^) and *P. vivax* case 2 (663.75 ± 5.39µm^2^; X^2^_2_ = 28.63, P <0.0001; Fig 3B). The average size of schizonts was negatively correlated to the sporozoite inoculum size (X^2^_7_ = 119.15, P <0.0001; Fig 3C). There was a significant interaction of *P. vivax* case and PHH lot (X^2^_6_ = 71.60, P <0.0001; Fig 3D) such that PHH lots providing the largest or smallest average schizont size were not always the same across infections with the three *P. vivax* cases.

**Fig 3.**
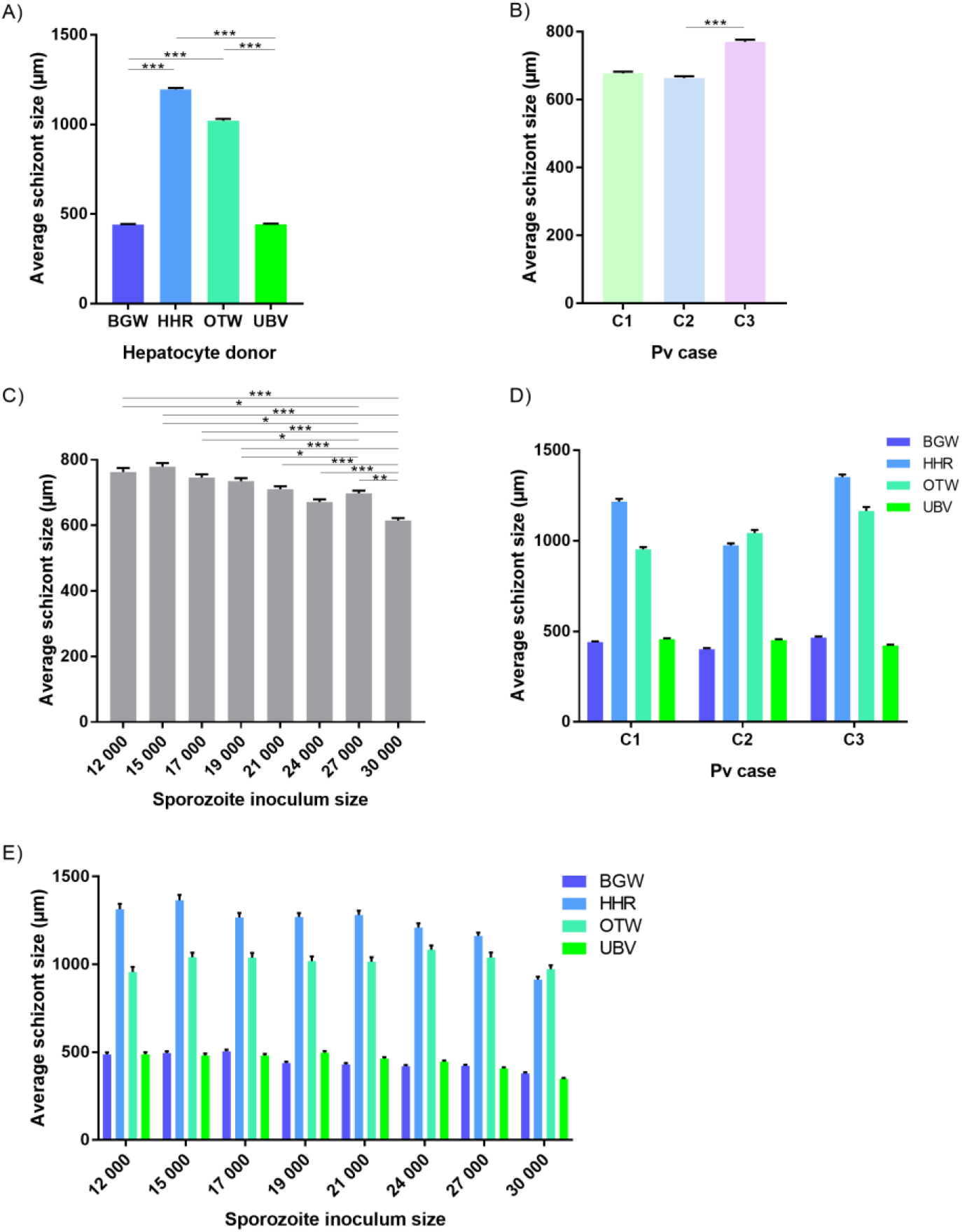
The average size of *P. vivax* schizonts by PHH lot and *P*. vivax case. Data include all parasites from Experiment 1 and are categorized by A) PHH lot, B) *P. vivax* isolate, C) sporozoite inoculum size, D) *P. vivax* isolate and PHH lot and E) sporozoite inoculum size and PHH lot. Asteriks indicate significant differences (Post hoc Tukey’s pairwise comparisons, *** P<0.0001, ** P<0.001, * P<0.05). Bars represent ± SE.

The differences in average schizont sizes between PHH lots tended to decrease as the sporozoite inoculum size increased; this effect was most apparent between lots HHR and OTW, which produced the largest schizonts of the four lots (PHH lots -sporozoite inoculum size interaction : X^2^_21_ = 42.84, P =0.0033; Fig 3E). The interaction between *P. vivax* case and sporozoite inoculum size was not significant (X^2^_14_ = 19.42, P = 0.15). Overall, UBV and BGW harbored a large proportion of small schizonts whereas HHR and OTW harbored similar proportions of schizonts of different size classes (S2 Fig). We did not investigate further a correlation between the total number of parasites in the well and the average schizont sizes as the data were segregated with UBV and BGW forming one group and OTW and HHR forming another group (S3 Fig).

### Experiment 2: comparisons of eight PHH donors and influence of sporozoite inoculum size

Eight different PHH lots were seeded and infected with 6 different sporozoite inoculums from a single *P. vivax* case. The average number of parasites per well was significantly influenced by the PHH lot (X^2^_7_ = 196.01, P <0.0001; Fig 4A) and the sporozoite inoculum size (X^2^_5_ = 192.24, P <0.0001; Fig 4B). The proportion of hypnozoites was significantly influenced by the PHH lot (X^2^_7_ =413.37, P <0.0001; Fig 4C) but not by the sporozoite inoculum size (X^2^_5_ = 11.026, P =0.051; Fig 4D).

**Fig 4.**
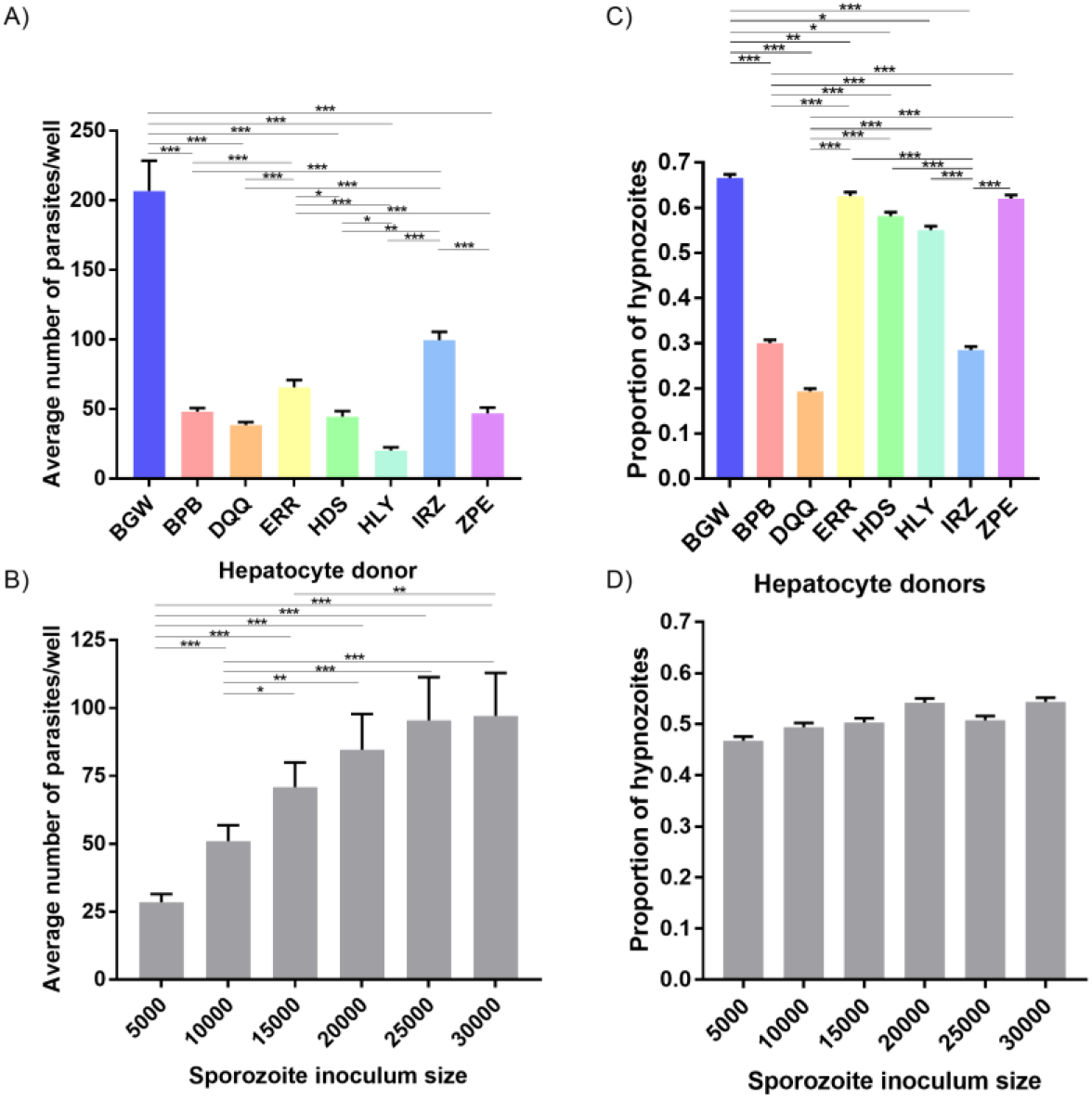
Growth metrics of liver stage parasites following infection of 8 PHH lots with one *P*.*vivax* case. Data include all parasites quantified from Experiment 2. The average number *P. vivax* parasites per well is shown categorized by A) PHH lot and B) sporozoite inoculum size. The average proportion of *P. vivax* hypnozoites per well is shown categorized by C) PHH lots and D) sporozoite inoculum size. Asteriks indicate significant differences (Post hoc Tukey’s pairwise comparisons, *** P<0.0001, ** P<0.001, * P<0.05). Bars represent ± SE (A,B) or ± 95% CI (C,D).

### Experiment 3: comparison of 51 *P. vivax* cases using one PHH donor

The total number of parasites per well was influenced by the *P. vivax* case (X^2^_50_ = 137.5, P <0.0001; Fig 5A) but not by the sporozoite inoculum size (X^2^ _1_=3.40, P=0.065). The average total number of parasites per well was influenced by the hepatocyte age at infection (X^2^_2_ =11.954, P=0.002). However, post-hoc comparisons showed only a significantly lower total number of parasites in hepatocytes infected at day 1 compared to day 2 post-seeding (81.83 ± 9.62 *vs*. 136.24 ± 7.12 respectively; Tukey’s post-hoc test: P=0.001, all other comparisons being non-significant). The average total number of parasites per well was not influenced by the presence of ABT (X^2^_1_ = 0.14, P=0.71), the season (X^2^ =0.41, P=0.52) nor the patient sex (X^2^ = 0.02, P=0.88). The proportion of hypnozoites was influenced by the *P. vivax* case (X^2^_50_ = 197.58, P <0.0001; Fig 5B) but not by the sporozoite inoculum size (X^2^_1_ = 0.69, P=0.40). The proportion of hypnozoites was also influenced by the hepatocyte age at infection (X^2^_2_=9.92, P=0.007). However post-hoc comparisons showed only a significantly lower proportion of hypnozoites in hepatocytes infected at day 3 compared to day 1 post-seeding (57.3 ± 0.7% vs. 63.3 ± 0.7%; Tukey’s post-hoc test: P=0.006, all other comparisons being non-significant). The proportion of hypnozoites was not affect by the sporozoite inoculum size (X^2^_1_ =0.69, P=0.40), the presence of ABT (X^2^ =2.99, P=0.08), the season (dry: 60.1 ± 0.7% vs. rainy: 66.6 ± 0.7%; X^2^_1_ =3.5, P=0.06) nor the patient sex (X^2^_1_ =0.10, P=0.75). The proportion of hypnozoites was not influenced by the number of visits to the clinic a patient already experienced (X^2^_1_ =1.36, P=0.24).

**Fig 5.**
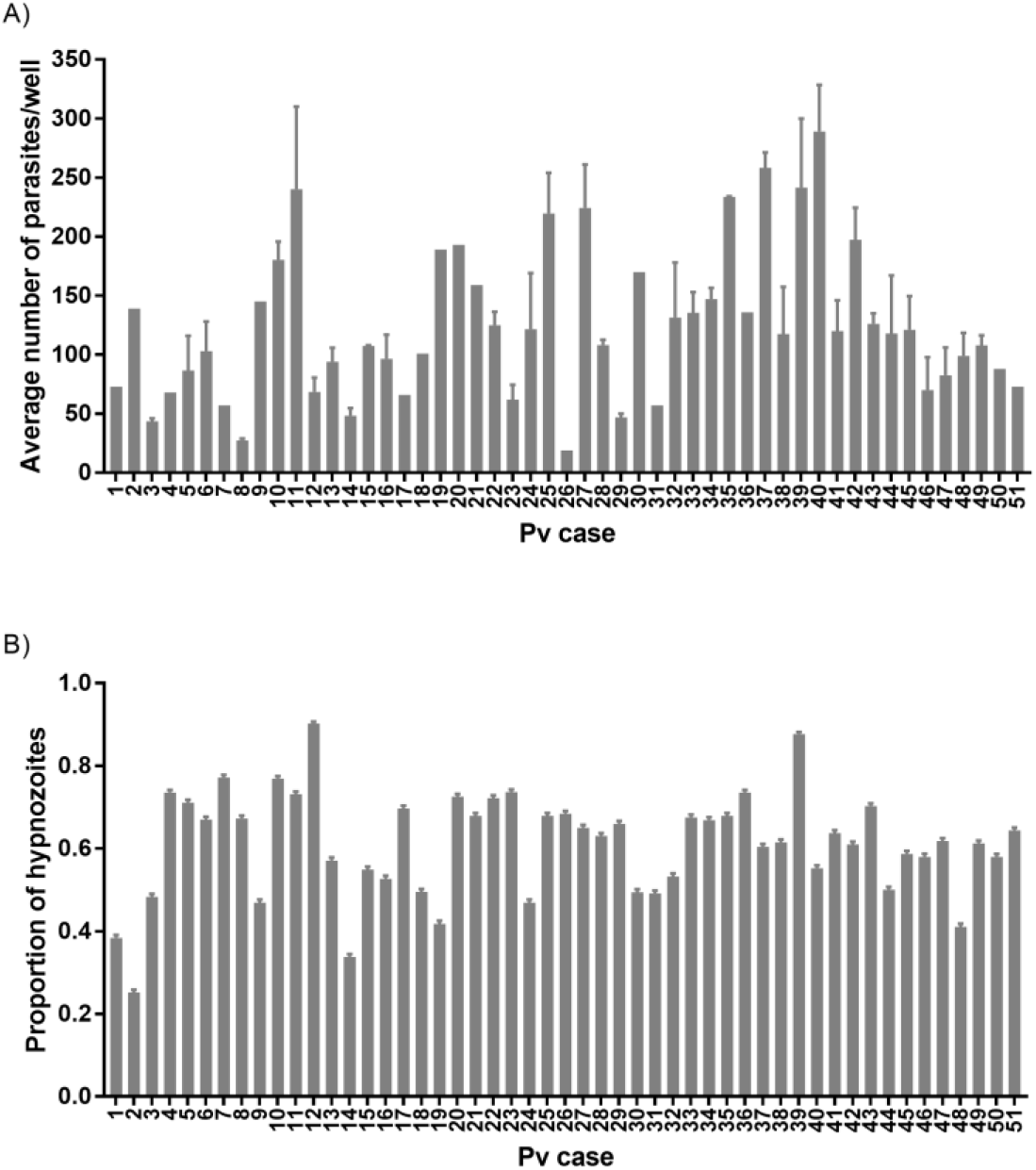
Growth metric of *P. vivax* liver stage parasites from 51 cases. A) Average number of parasites per well and B) proportion of hypnozoites. Asteriks indicate significant differences (Post hoc Tukey’s pairwise comparisons, *** P<0.0001, ** P<0.001, * P<0.05). Bars represent ± 95% CI.

## Discussion

The determinants of relapse periodicity and the molecular processes that drive hypnozoite formation, persistence and activation are still largely unknown. Three non-mutually exclusive hypotheses have been proposed regarding the determinants of hypnozoite formation: predetermination during the sporozoite development in the mosquito, fate determination as sporozoites progress from mosquito salivary glands to the hepatocytes, and fate determination after infecting the host hepatocyte [reviewed in 33]. We found that both the PHH donor lot and the *P. vivax* case influenced the proportion of hypnozoites observed (Figs 2, 4). This suggests that the proportion of hypnozoites is strain-dependent as previously shown in a humanized mouse model [15]. Although laboratory conditions have some variability, the large differences in hypnozoite proportions observed with infections from 51 *P. vivax* cases using one hepatocyte donor over time further support this hypothesis. Interestingly, a recent report suggests sporozoites found in mosquito salivary glands do not seem to present two distinct transcriptional signatures corresponding to future hypnozoites and schizonts[34], further suggesting that transcriptional changes responsible for the fate of sporozoites could occur in the host hepatocyte or be epigenetically-controlled [33].

The different hepatocyte donors we tested showed large variation in both the quantity and ratio of hypnozoites to schizonts after infection. We performed two experiments comparing multiple donors; one with four donors previously identified as highly supportive of *P. vivax* infection (Fig 1A), and another with seven donors supporting various levels of *P. vivax* infection in comparison to the highly-supportive lot BGW (Fig 4A). These results suggest host hepatocytes harbor characteristics making them more-or less-supportive of a quiescent versus growing intracellular parasite. Indeed, recent studies have shown that liver stage parasites must form and maintain a delicate interface with the host hepatocyte’s lysosomes, with failure leading to parasite death [35]. This survivability factor is one example of many possible host-parasite interactions that could explain both the net parasite and ratio differences we noted across donors. Another explanation for the infection rate differences is that, while sporozoites are traversing hepatocytes, there may sense the suitability of a particular hepatocyte prior to switching from traversal to invasion, which is distinctly marked by formation of a parasitophorous vacuole membrane (reviewed in [36]). Such a tropism has been described for *P. yoelii* and *P. falciparum* sporozoites preferentially infecting polyploid hepatocytes [37]. Regarding the fate of sporozoites in these different donor lots, it has been shown that *P. vivax* hypnozoites are susceptible to several antimalarials for the first few days post-hepatocyte infection [38, 39]. During this time, the parasite’s cytoplasmic compartment and membranes grow to several times the volume of a sporozoite and begin to incorporate host proteins such as aquaporin 3, indicating sporozoites must establish quiescence, and are not immediately quiescent, following infection [39]. Likewise, recent reports of the earliest known markers of liver schizogony, including DNA synthesis, division of the parasite nucleus, and expression of liver-specific protein 2 at 3 days post-hepatocyte infection, suggest that commitment to schizogony may not occur immediately after hepatocyte infection [15, 40, 41]. As there seems to be ample time for a cell-cycle checkpoint to prevent DNA synthesis as liver forms are established, we speculate that, over the first 24-48 hours post-infection, sporozoites may be able to sense, or at least be influenced by, the intracellular environment of the hepatocyte, and then respond to specific conditions or stimuli by forming either a hypnozoite or schizont. Yet another possible explanation for the different infection rate and hypnozoite ratio noted across donors could be the composition of hepatocytes in each lot as from either zone 1, 2, or 3 of the liver lobule. Liver lobules perform specific metabolic functions and display different levels of glycolysis and cellular respiration [20-22]. Recently, a study of primary hepatocytes infected with *P. falciparum* demonstrated cultured primary human hepatocytes are comprised of different ratios of cells from each zone and zonal differentiation as important for liver stage development. In our recent report of single-cell RNA-sequencing of *P. vivax* liver stages and host hepatocytes, we did look for an infection and fating preference for sporozoites infected into a culture of various hepatocyte subpopulations, however we found no clear infection pattern or preference [23]. Further deciphering the components of the host cell environment as allowing or favoring hypnozoites versus schizont formation would help better understand the mechanisms of dormancy.

In Cambodia, vivax malaria is less-frequently transmitted during the dry season, when the population of the Anopheline vector is at its lowest level. During this time, it would be advantageous for any vivax parasites that do transmit to form hypnozoites, such that the blood stage infections resulting from transmission would occur after the end of the dry season. Such a mechanism has been described for strains of long-latency vivax malaria such as those formerly prevalent in northern latitudes [42]. However, seasonal variation in Cambodia might not be strong enough to select for long-latency strains. It has also been shown that, for some strains, once relapses begin after a period of latency, they are frequent, indicating cessation of latency is also be programmed [6]. In this study, we had the unique opportunity to quantify the formation of hypnozoites and schizonts of *P. vivax* isolates from patients during the dry season as well as from the same patients visiting the clinic for malaria therapies multiple times. While we did not find an apparent effect of the same patient recurrently visiting the clinic nor of the season on the proportion of hypnozoites, we did find that some of the 51 cases exhibited remarkably high hypnozoite ratios, indicating genotypes encoding for hypnozoite formation do persist in the population and likely factor into latency and ongoing transmission. As we were not able to collect enough parasite material for DNA or RNA sequencing to further characterize these unique cases, future studies could combine donor panels with a multi-omics approach to better understand these genotype-phenotype relationships.

A crowding effect could influence sporozoites to become hypnozoites to avoid competition and increase the chance of transmission by opportunistically causing a relapse after the primary blood infection and subsequent immune response. To look for such a crowding effect on sporozoite fating we performed two experiments with an inoculum gradient culminating with a highest inoculum of 30 ×10^3^ sporozoites per well, or a relatively large multiplicity of infection of >2 sporozoites for each primary hepatocyte. As expected, the number of parasites increased positively with the sporozoite inoculum size in these two first experiments, resulting in an apparent plateau representing saturation (Fig 1C, Fig 4B), However, we observed only a small influence of the sporozoite inoculum size on the proportion of hypnozoites in only one experiment out of three which suggest that this factor is not likely a strong determinant of sporozoite fating. In our third experiment we further analyzed the effect of inoculum size and found it did not influence the infection rate of 51 *P. vivax* cases used for a routine drug screening program. Thus, the sporozoite inoculum size influences the infection rate with some modulations, which could be due to an intrinsic *P. vivax* case difference in infectivity or to the sporozoite development status. Indeed, salivary gland dissections result in the collection of all sporozoite available, not only the mature sporozoites which would have migrated within the saliva during a natural mosquito-bite infection. Therefore, developmental heterogeneity of sporozoites could explain the different infection rates observed across the *P. vivax* cases used [34, 43]. Comparing hypnozoite ratios in hepatocyte infections resulting from dissections of one batch of infected mosquitoes over several days would help resolve an effect of sporozoite loiter time in vector.

We found that schizont growth was strongly influenced by the hepatocyte donor. Interestingly, two patterns were observed in *P. vivax* infections independent from a parasite density effect: hepatocyte donors supporting a large proportion of small schizonts and few large schizonts versus hepatocyte donors supporting a homogenous distribution of schizont sizes. Comparing the net production of merozoites would be interesting to determine if the two different strategies results in different parasite loads. Similarly, comparing the individual cell metabolic activities of these two types of donor would help our understanding of the factors driving schizont development.

In conclusion, elegant studies have shown the first relapses in life are genetically homologous and that the parasites causing relapses in a vivax malaria patient were likely caused by hypnozoites from meiotic sibling sporozoites from the oocyst phase of the lifecycle [14, 44]. These studies suggest genetic crosses could be used to further investigate the determinates of sporozoite fating under controlled laboratory conditions. Such studies would be remarkably informative if methods are ever established for *in vitro* propagation and experimental transmission of *P. vivax* strains. In lieu of such an experimental system, *P. vivax* strains with distinct relapse phenotypes can be propagated and transmitted from experimental infections of nonhuman primates [reviewed in 45], or perhaps humanized mice [46]. These systems would allow interrogation of *P. vivax* sporozoite fating with either wild-type or transgenic parasites strains and do so with experimental replication with parasites of the same genetic background.

This report utilizes a well-controlled experimental design to identify and measure the relative effect of factors influencing sporozoite invasion and development in a hepatocyte culture system and, having used patient isolates to generate sporozoites, serves as a natural reference point as investigators focus on understanding hypnozoite biology using these alternative model systems.

## Acknowledgments

We thank the patients of Mondulkiri Province, Cambodia, for participating in this study. High content imaging data was obtain in the Biomedical Microscopy Core at the University of Georgia, supported by the Georgia Research Alliance.

## Author contributions

Conceptualization: A.V., J.P., S.M.; Data curation, formal analysis, visualization: A. V.; Funding Acquisition, Project Administration and Resources : A.V., B. W., S. M., D. K.; Investigation: J.P., C.C., A.V., S.M.; Writing of original draft and preparation: A.V. and S.M. All co-authors reviewed, edited and approved the manuscript.

## Data availability

All data files will be available from the Dryad database.

## Supporting Information

**S1 Fig. Plate map of *P. vivax* experiment 1**.

**S2 Fig. Proportion of schizonts in each size class for each PHH lot, *P. vivax* case, and sporozoite inoculum size**.

**S3 Fig. Plot of the averaged schizont size by the number of parasites per well**.

