## Supplementary material for "Liver stage fate determination in *Plasmodium vivax* parasites: characterization of schizont growth and hypnozoite fating from patient isolates": figure S1

### Plate maps

#### Plate 1

<table>
<tr><th></th><th colspan="12">UBV</th><th colspan="12">HHR</th></tr>
<tr><th></th><th>1</th><th>2</th><th>3</th><th>4</th><th>5</th><th>6</th><th>7</th><th>8</th><th>9</th><th>10</th><th>11</th><th>12</th><th>13</th><th>14</th><th>15</th><th>16</th><th>17</th><th>18</th><th>19</th><th>20</th><th>21</th><th>22</th><th>23</th><th>24</th></tr>
<tr><td>A</td><td></td><td></td><td></td><td></td><td></td><td></td><td></td><td></td><td></td><td></td><td></td><td></td><td></td><td></td><td></td><td></td><td></td><td></td><td></td><td></td><td></td><td></td><td></td><td></td></tr>
<tr><td>B</td><td></td><td rowspan="6">6000</td><td rowspan="6">9000</td><td rowspan="6">12000</td><td rowspan="6">15000</td><td rowspan="6">17000</td><td rowspan="6">19000</td><td rowspan="6">21000</td><td rowspan="6">25000</td><td rowspan="6">27000</td><td rowspan="6">30000</td><td></td><td></td><td rowspan="6">6000</td><td rowspan="6">9000</td><td rowspan="6">12000</td><td rowspan="6">15000</td><td rowspan="6">17000</td><td rowspan="6">19000</td><td rowspan="6">21000</td><td rowspan="6">25000</td><td rowspan="6">27000</td><td rowspan="6">30000</td><td></td></tr>
<tr><td>C</td><td></td><td></td><td></td><td></td></tr>
<tr><td>D</td><td></td><td></td><td></td><td></td></tr>
<tr><td>E</td><td></td><td></td><td></td><td></td></tr>
<tr><td>F</td><td></td><td></td><td></td><td></td></tr>
<tr><td>G</td><td></td><td></td><td></td><td></td></tr>
<tr><td>H</td><td></td><td></td><td></td><td></td><td></td><td></td><td></td><td></td><td></td><td></td><td></td><td></td><td></td><td></td><td></td><td></td><td></td><td></td><td></td><td></td><td></td><td></td><td></td><td></td></tr>
</table>

**Pv case 1**

#### Plate 2

<table>
<tr><th></th><th colspan="12">BGW</th><th colspan="12">OTW</th></tr>
<tr><th></th><th>1</th><th>2</th><th>3</th><th>4</th><th>5</th><th>6</th><th>7</th><th>8</th><th>9</th><th>10</th><th>11</th><th>12</th><th>13</th><th>14</th><th>15</th><th>16</th><th>17</th><th>18</th><th>19</th><th>20</th><th>21</th><th>22</th><th>23</th><th>24</th></tr>
<tr><td>A</td><td></td><td></td><td></td><td></td><td></td><td></td><td></td><td></td><td></td><td></td><td></td><td></td><td></td><td></td><td></td><td></td><td></td><td></td><td></td><td></td><td></td><td></td><td></td><td></td></tr>
<tr><td>B</td><td></td><td rowspan="6">6000</td><td rowspan="6">9000</td><td rowspan="6">12000</td><td rowspan="6">15000</td><td rowspan="6">17000</td><td rowspan="6">19000</td><td rowspan="6">21000</td><td rowspan="6">25000</td><td rowspan="6">27000</td><td rowspan="6">30000</td><td></td><td></td><td rowspan="6">6000</td><td rowspan="6">9000</td><td rowspan="6">12000</td><td rowspan="6">15000</td><td rowspan="6">17000</td><td rowspan="6">19000</td><td rowspan="6">21000</td><td rowspan="6">25000</td><td rowspan="6">27000</td><td rowspan="6">30000</td><td></td></tr>
<tr><td>C</td><td></td><td></td><td></td><td></td></tr>
<tr><td>D</td><td></td><td></td><td></td><td></td></tr>
<tr><td>E</td><td></td><td></td><td></td><td></td></tr>
<tr><td>F</td><td></td><td></td><td></td><td></td></tr>
<tr><td>G</td><td></td><td></td><td></td><td></td></tr>
<tr><td>H</td><td></td><td></td><td></td><td></td><td></td><td></td><td></td><td></td><td></td><td></td><td></td><td></td><td></td><td></td><td></td><td></td><td></td><td></td><td></td><td></td><td></td><td></td><td></td><td></td></tr>
</table>

**Pv case 1**

#### Plate 3

<table>
<tr><th></th><th colspan="12">UBV</th><th colspan="12">HHR</th></tr>
<tr><th></th><th>1</th><th>2</th><th>3</th><th>4</th><th>5</th><th>6</th><th>7</th><th>8</th><th>9</th><th>10</th><th>11</th><th>12</th><th>13</th><th>14</th><th>15</th><th>16</th><th>17</th><th>18</th><th>19</th><th>20</th><th>21</th><th>22</th><th>23</th><th>24</th></tr>
<tr><td>A</td><td></td><td></td><td></td><td></td><td></td><td></td><td></td><td></td><td></td><td></td><td></td><td></td><td></td><td></td><td></td><td></td><td></td><td></td><td></td><td></td><td></td><td></td><td></td><td></td></tr>
<tr><td>B</td><td></td><td></td><td></td><td rowspan="6">12000</td><td rowspan="6">15000</td><td rowspan="6">17000</td><td rowspan="6">19000</td><td rowspan="6">21000</td><td rowspan="6">25000</td><td rowspan="6">27000</td><td rowspan="6">30000</td><td></td><td></td><td></td><td></td><td rowspan="6">12000</td><td rowspan="6">15000</td><td rowspan="6">17000</td><td rowspan="6">19000</td><td rowspan="6">21000</td><td rowspan="6">25000</td><td rowspan="6">27000</td><td rowspan="6">30000</td><td></td></tr>
<tr><td>C</td><td></td><td></td><td></td><td></td><td></td><td></td><td></td><td></td></tr>
<tr><td>D</td><td></td><td></td><td></td><td></td><td></td><td></td><td></td><td></td></tr>
<tr><td>E</td><td></td><td></td><td></td><td></td><td></td><td></td><td></td><td></td></tr>
<tr><td>F</td><td></td><td></td><td></td><td></td><td></td><td></td><td></td><td></td></tr>
<tr><td>G</td><td></td><td></td><td></td><td></td><td></td><td></td><td></td><td></td></tr>
<tr><td>H</td><td></td><td></td><td></td><td></td><td></td><td></td><td></td><td></td><td></td><td></td><td></td><td></td><td></td><td></td><td></td><td></td><td></td><td></td><td></td><td></td><td></td><td></td><td></td><td></td></tr>
<tr><td>I</td><td></td><td></td><td></td><td></td><td></td><td></td><td></td><td></td><td></td><td></td><td></td><td></td><td></td><td></td><td></td><td></td><td></td><td></td><td></td><td></td><td></td><td></td><td></td><td></td></tr>
<tr><td>J</td><td></td><td rowspan="6">6000</td><td rowspan="6">9000</td><td rowspan="6">12000</td><td rowspan="6">15000</td><td rowspan="6">17000</td><td rowspan="6">19000</td><td rowspan="6">21000</td><td rowspan="6">25000</td><td rowspan="6">27000</td><td rowspan="6">30000</td><td></td><td></td><td rowspan="6">6000</td><td rowspan="6">9000</td><td rowspan="6">12000</td><td rowspan="6">15000</td><td rowspan="6">17000</td><td rowspan="6">19000</td><td rowspan="6">21000</td><td rowspan="6">25000</td><td rowspan="6">27000</td><td rowspan="6">30000</td><td></td></tr>
<tr><td>K</td><td></td><td></td><td></td><td></td></tr>
<tr><td>L</td><td></td><td></td><td></td><td></td></tr>
<tr><td>M</td><td></td><td></td><td></td><td></td></tr>
<tr><td>N</td><td></td><td></td><td></td><td></td></tr>
<tr><td>O</td><td></td><td></td><td></td><td></td></tr>
<tr><td>P</td><td></td><td></td><td></td><td></td><td></td><td></td><td></td><td></td><td></td><td></td><td></td><td></td><td></td><td></td><td></td><td></td><td></td><td></td><td></td><td></td><td></td><td></td><td></td><td></td></tr>
</table>

**Pv case 2**

**Pv case 3**

#### Plate 4

<table>
<tr><th></th><th colspan="12">BGW</th><th colspan="12">OTW</th></tr>
<tr><th></th><th>1</th><th>2</th><th>3</th><th>4</th><th>5</th><th>6</th><th>7</th><th>8</th><th>9</th><th>10</th><th>11</th><th>12</th><th>13</th><th>14</th><th>15</th><th>16</th><th>17</th><th>18</th><th>19</th><th>20</th><th>21</th><th>22</th><th>23</th><th>24</th></tr>
<tr><td>A</td><td></td><td></td><td></td><td></td><td></td><td></td><td></td><td></td><td></td><td></td><td></td><td></td><td></td><td></td><td></td><td></td><td></td><td></td><td></td><td></td><td></td><td></td><td></td><td></td></tr>
<tr><td>B</td><td></td><td></td><td></td><td rowspan="6">12000</td><td rowspan="6">15000</td><td rowspan="6">17000</td><td rowspan="6">19000</td><td rowspan="6">21000</td><td rowspan="6">25000</td><td rowspan="6">27000</td><td rowspan="6">30000</td><td></td><td></td><td></td><td></td><td rowspan="6">12000</td><td rowspan="6">15000</td><td rowspan="6">17000</td><td rowspan="6">19000</td><td rowspan="6">21000</td><td rowspan="6">25000</td><td rowspan="6">27000</td><td rowspan="6">30000</td><td></td></tr>
<tr><td>C</td><td></td><td></td><td></td><td></td><td></td><td></td><td></td><td></td></tr>
<tr><td>D</td><td></td><td></td><td></td><td></td><td></td><td></td><td></td><td></td></tr>
<tr><td>E</td><td></td><td></td><td></td><td></td><td></td><td></td><td></td><td></td></tr>
<tr><td>F</td><td></td><td></td><td></td><td></td><td></td><td></td><td></td><td></td></tr>
<tr><td>G</td><td></td><td></td><td></td><td></td><td></td><td></td><td></td><td></td></tr>
<tr><td>H</td><td></td><td></td><td></td><td></td><td></td><td></td><td></td><td></td><td></td><td></td><td></td><td></td><td></td><td></td><td></td><td></td><td></td><td></td><td></td><td></td><td></td><td></td><td></td><td></td></tr>
<tr><td>I</td><td></td><td></td><td></td><td></td><td></td><td></td><td></td><td></td><td></td><td></td><td></td><td></td><td></td><td></td><td></td><td></td><td></td><td></td><td></td><td></td><td></td><td></td><td></td><td></td></tr>
<tr><td>J</td><td></td><td rowspan="6">6000</td><td rowspan="6">9000</td><td rowspan="6">12000</td><td rowspan="6">15000</td><td rowspan="6">17000</td><td rowspan="6">19000</td><td rowspan="6">21000</td><td rowspan="6">25000</td><td rowspan="6">27000</td><td rowspan="6">30000</td><td></td><td></td><td rowspan="6">6000</td><td rowspan="6">9000</td><td rowspan="6">12000</td><td rowspan="6">15000</td><td rowspan="6">17000</td><td rowspan="6">19000</td><td rowspan="6">21000</td><td rowspan="6">25000</td><td rowspan="6">27000</td><td rowspan="6">30000</td><td></td></tr>
<tr><td>K</td><td></td><td></td><td></td><td></td></tr>
<tr><td>L</td><td></td><td></td><td></td><td></td></tr>
<tr><td>M</td><td></td><td></td><td></td><td></td></tr>
<tr><td>N</td><td></td><td></td><td></td><td></td></tr>
<tr><td>O</td><td></td><td></td><td></td><td></td></tr>
<tr><td>P</td><td></td><td></td><td></td><td></td><td></td><td></td><td></td><td></td><td></td><td></td><td></td><td></td><td></td><td></td><td></td><td></td><td></td><td></td><td></td><td></td><td></td><td></td><td></td><td></td></tr>
</table>

**Pv case 2**

**Pv case 3**
