## Supplementary figures and images for "Liver stage fate determination in *Plasmodium vivax* parasites: characterization of schizont growth and hypnozoite fating from patient isolates"

### figure S2

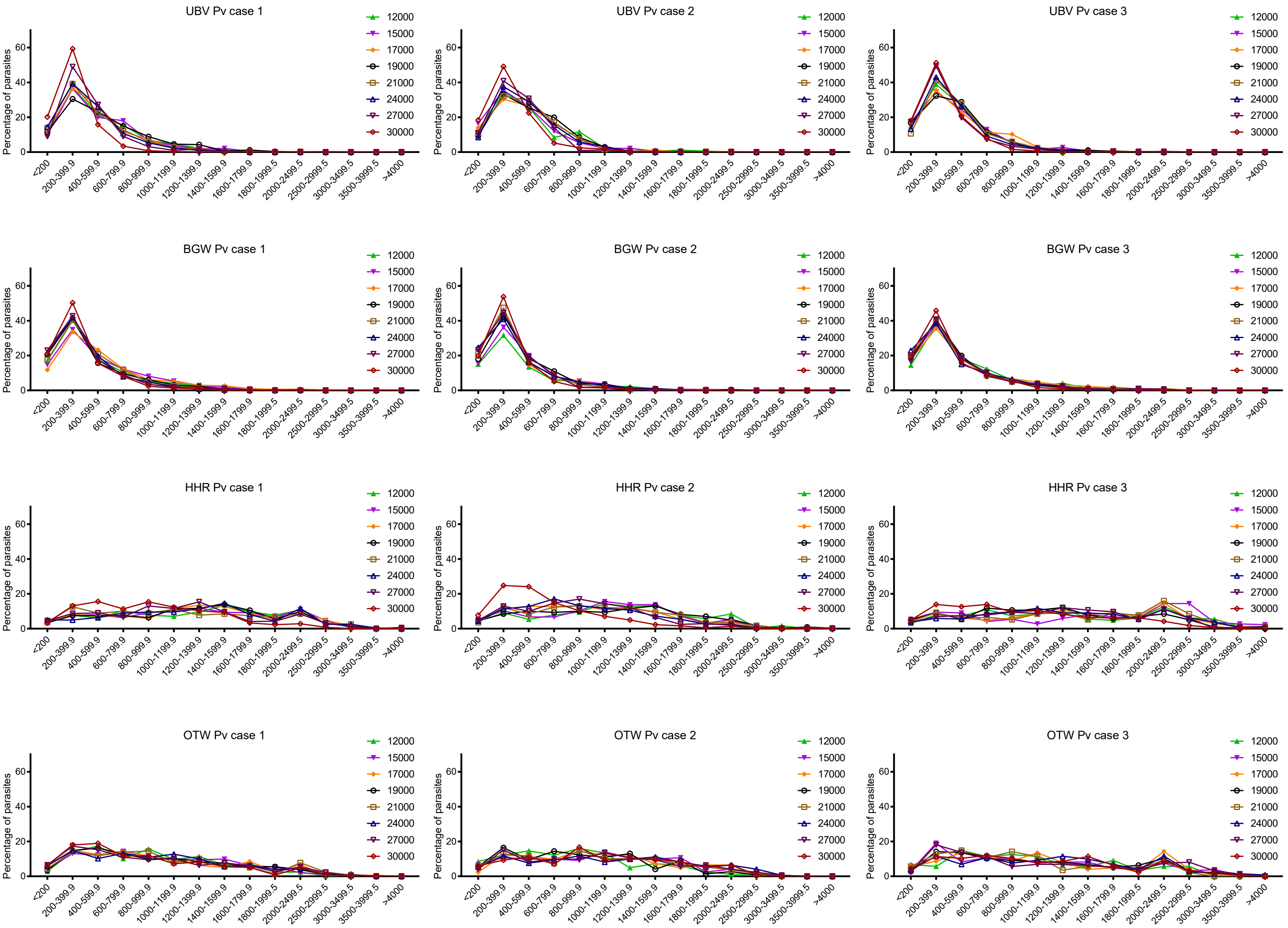

### figure S3

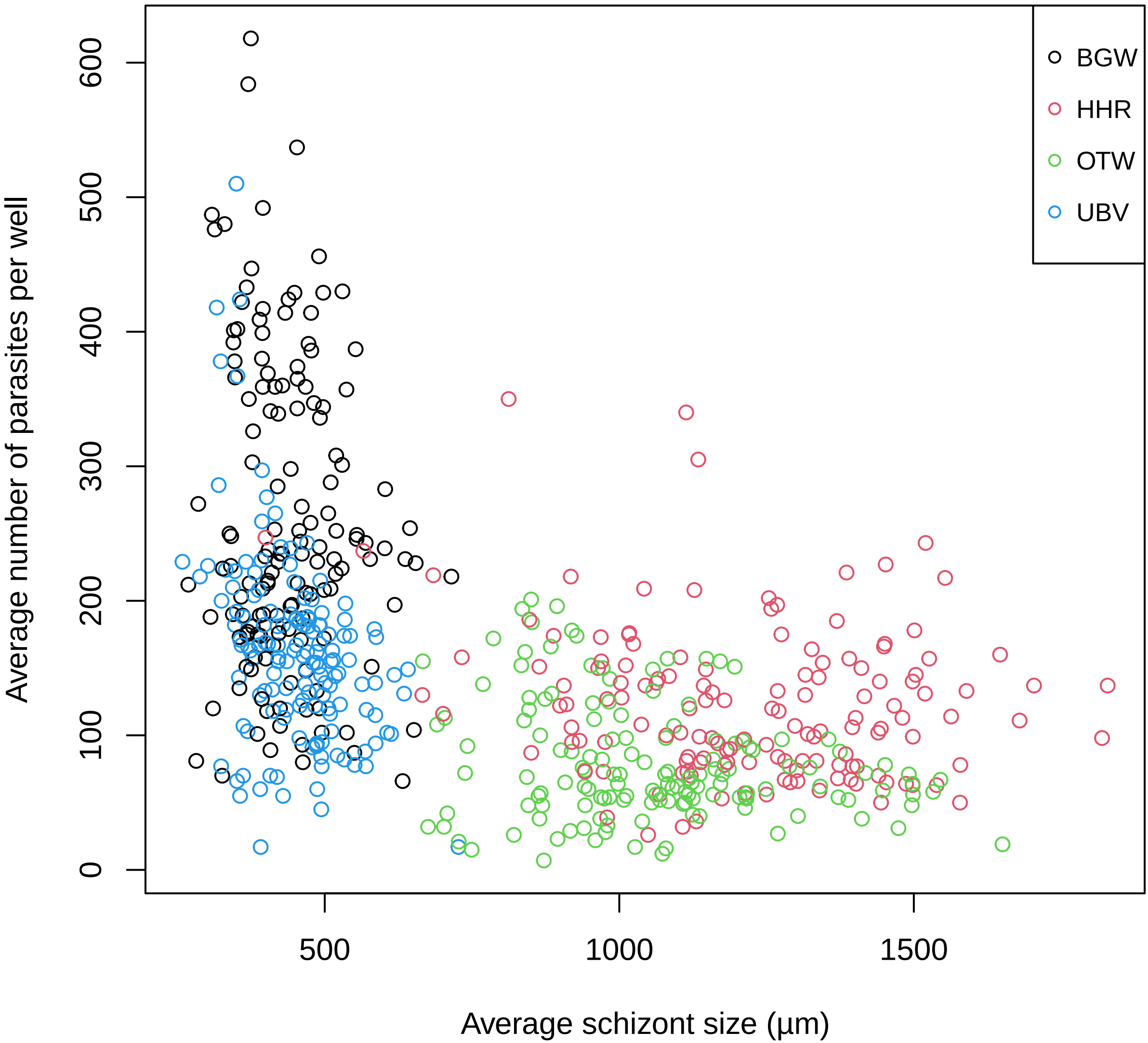
